# The voltage-gated sodium channel Na_V_1.7 underlies endometriosis-associated chronic pelvic pain

**DOI:** 10.1101/2022.10.06.511228

**Authors:** Joel Castro, Jessica Maddern, Chuen Yuen Chow, Poanna Tran, Irina Vetter, Glenn F. King, Stuart M. Brierley

**Author notes:** Corresponding author: Dr Joel Castro (PhD), Visceral Pain Research Group, Level 7, South Australian Health and Medical Research Institute (SAHMRI), North Terrace, SA 5000, Australia.

## Abstract

Chronic pelvic pain (CPP) is the primary symptom of endometriosis patients, but adequate treatments are lacking. Modulation of ion channels expressed by sensory nerves innervating the viscera have shown promise for the treatment of irritable bowel syndrome and overactive bladder. However, similar therapies have not been explored for endometriosis-associated CPP. Here we examined the role of the voltage-gated sodium (Na_V_) channel Na_V_1.7 in the sensitivity of vagina-innervating sensory afferents and investigated whether Na_V_1.7 inhibition reduces nociceptive signals from the vagina and ameliorates endometriosis-associated CPP. The mechanical responsiveness of vagina-innervating sensory afferents was assessed with *ex vivo* single unit recording preparations. Pain evoked by vaginal distension (VD) was quantified by the visceromotor response (VMR) *in vivo*. In control mice, pharmacological activation of Na_V_1.7 with OD1 sensitised vagina-innervating pelvic afferents to mechanical stimuli. Using a syngeneic mouse model of endometriosis, we established that endometriosis sensitized vagina-innervating pelvic afferents to mechanical stimuli. The highly selective Na_V_1.7 inhibitor Tsp1a revealed that this afferent hypersensitivity occurred in a Na_V_1.7-dependent manner. Moreover, *in vivo* intra-vaginal treatment with Tsp1a reduced the exaggerated VMRs to VD that is characteristic of mice with endometriosis. Conversely, Tsp1a did not alter *ex vivo* afferent mechanosensitivity or *in vivo* VMRs to VD in Sham control mice. Collectively, these findings suggest that Na_V_1.7 plays a crucial role in endometriosis-induced vaginal hyperalgesia. Importantly, Na_V_1.7 inhibition selectively alleviated endometriosis-associated CPP without the loss of normal sensation, suggesting that selective targeting of Na_V_1.7 could improve the quality of life of women with endometriosis.

## Introduction

Endometriosis is a debilitating condition characterised by infertility and chronic pelvic pain (CPP) that affects approximately 11% of women worldwide [4; 25]. Vaginal hyperalgesia, also known as dyspareunia, or painful intercourse, is one of the most debilitating symptoms affecting women with endometriosis. Current treatments aimed at relieving the CPP associated with endometriosis include hormonal suppression of ovarian function or surgery. Unfortunately, both interventions have variable success and negatively impact fertility [61], reflecting a clear need for alternative treatment options.

Physiological and painful stimuli are detected by ion channels and receptors expressed within the terminals of sensory afferent nerve fibres projecting from the periphery to the central nervous system (CNS). Modulation of ion channels expressed within sensory neurons innervating the colon and the bladder has proved be a promising approach for treatment of chronic pain associated with irritable bowel syndrome (IBS) and overactive bladder syndrome (OAB) [9; 32; 40; 41; 51; 55; 56]. Remarkably, a similar approach directed to relieve endometriosis-associated CPP has yet to be described. This is partially due to the lack of knowledge of specific ion channels and receptors expressed within sensory neurons projecting to the female reproductive tract. In fact, only the transient receptor potential vanilloid 1 channel TRPV1 [20], purinergic receptor P2X_3_ [59], voltage-gated sodium (Na_V_) channels [16; 45], and the voltage-gated potassium channels (K_v_) K_V_6.4 and K_V_2.1 [45], have been reported to be expressed in these sensory afferents. While only tetrodotoxin (TTX)-sensitive Na_V_ channels have been reported to have a functional role in pain sensation from the female reproductive tract of healthy mice [16], the identity of the Na_V_ channel(s) contributing to this pain signalling pathway, and whether they contribute to endometriosis-associated CPP, remains unknown.

The Na_V_ channel family consists of nine isoforms (Na_V_1.1–Na_V_1.9) that are characterised as either TTX-sensitive (Na_V_1.1–Na_V_1.4, Na_V_1.6, and Na_V_1.7) or TTX-resistant (Na_V_1.5, Na_V_1.8, and Na_V_1.9) [18; 19]. Na_V_1.7 is a therapeutic target of interest as loss-of-function mutations in the gene encoding Na_V_1.7 (*SCN9A*) lead to a congenital inability to sense pain, while gain-of-function mutations lead to increased pain perception [5; 42]. Some Na_V_1.7 inhibitors have been tested in clinical trials for different types of pain [1; 3; 29; 44; 60]. Despite the abundant expression of Na_V_1.7 within visceral-innervating neurons (98–100% expression in colon-, bladder- and vagina-innervating lumbosacral DRG neurons [16; 32; 38], the contribution of Na_V_1.7 to visceral sensation and pain is controversial [26]. Some studies suggest that Na_V_1.7 is involved in somatic, but not visceral, pathways in acute pain [36], while others have shown that it plays an important role in the chronic abdominal pain associated with IBS [41].

In this study we used a highly selective inhibitor of Na_V_1.7 [41] to investigate the role of this channel in vaginal sensation and evoked pain. Additionally, we examined whether Na_V_1.7 contributes to the allodynia and hyperalgesia experienced by mice with endometriosis. We show that Na_V_1.7 contributes to endometriosis-associated vaginal hyperalgesia, which can be ameliorated by treatment with a Nav1.7 inhibitor. Our data suggest that inhibition of Na_V_1.7 may be a viable option to treat endometriosis-associated CPP.

## Results

### Activation of Na_V_1.7 increases vaginal afferent sensitivity to mechanical stimuli

Little is known about how mechanical stimuli is sensed by female reproductive organs. We recently showed that the sensitivity of pelvic vaginal afferents to mechanical stimuli can be altered by nonspecific modulators of Na_V_ channels [16]. This includes enhancing afferent responses to mechanical stimuli with veratridine and conversely inhibiting responses with TTX [16]. In this study, we dissected these mechanisms further by investigating the specific contribution of Na_V_1.7, which is expressed by 100% of vagina-innervating neurons [16].

Compared to baseline responses, application of the α-scorpion toxin OD1, a selective agonist of Na_V_1.7 [23], significantly increased pelvic vaginal afferent responses to vaginal stroking (**Figure 1Ai-iii**), focal compression (**Figure 1Bi-ii**), and circular stretch (**Figure 1Ci-ii**). In addition to an increased number of action potentials (AP) generated by circular stretch, OD1 significatively reduced the latency of the response (time for the first AP to be generated) to circular stretch (**Figure 1Ciii**). Interestingly, greater than 50 % of the vaginal afferents studied continued to fire APs after cessation of the mechanical stimuli (**Figure 1Ci**). Overall, these results indicate that the mechanical responsiveness of vaginal afferents can be augmented by activation of Na_V_1.7 expressed within these fibres.

**Figure 1:**
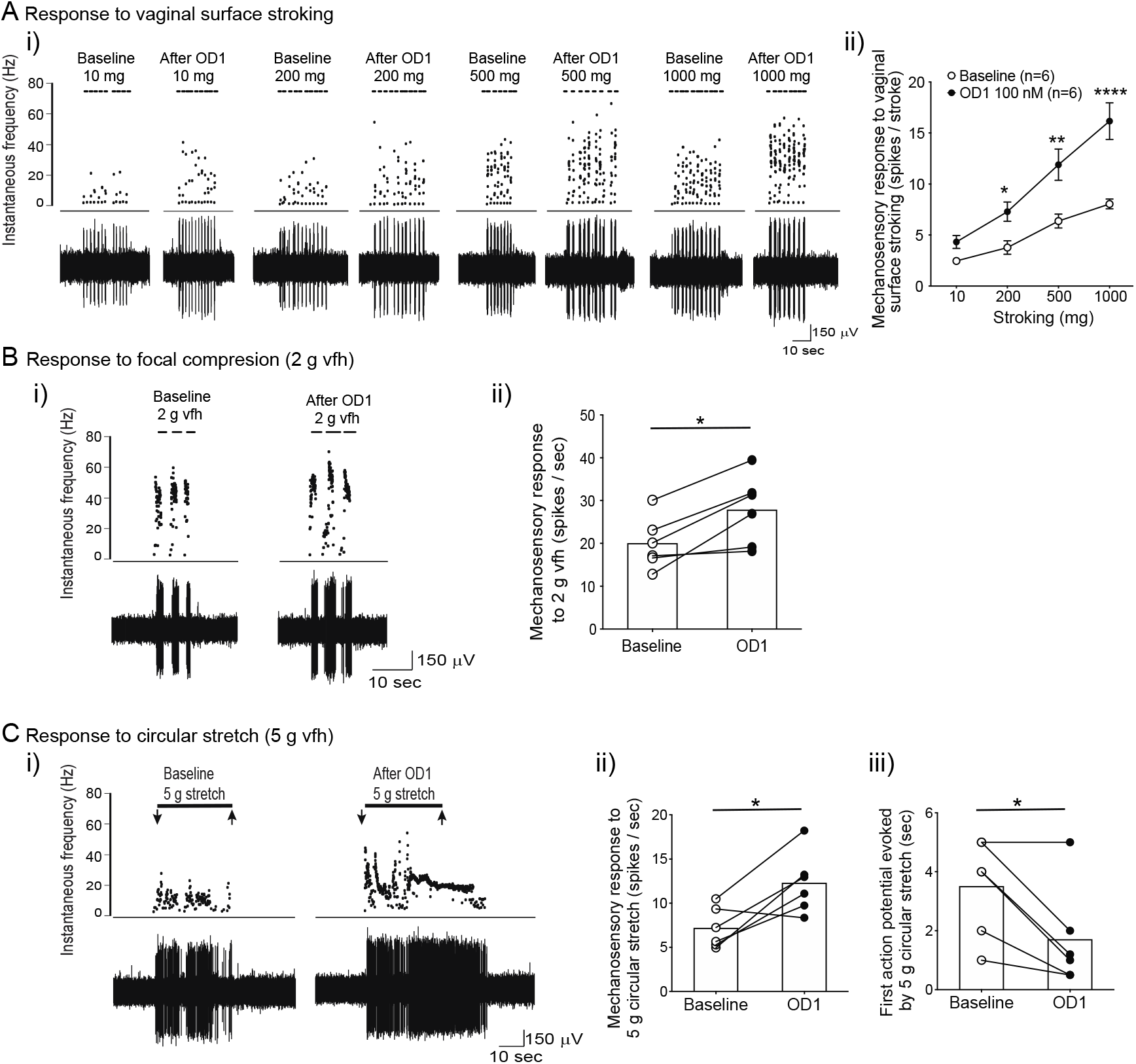
Pelvic vaginal afferents from Sham control mice display enhanced responses to a variety of mechanical stimuli following activation of Na_V_1.7 with OD1. **(A)** (**i)** Representative traces obtained with an ex vivo single-unit nerve recording preparation, showing AP discharge of vaginal afferents in response to graded stroking of the vagina surface at baseline and following incubation of the receptive field with of OD1 (100 nM) for 5 min. (**ii)** Group data showing that OD1 significatively increased AP discharge elicited by stroking of the vagina surface (*P<0.05 at 200 mg, **P<0.01 at 500 mg, and ***P<0.0001 at 1000 mg, two-way ANOVA followed by Bonferroni post-hoc comparison tests). (**B)** (**i)** Representative traces of vaginal afferent responses to focal compression with a 2 g vfh filament at baseline and following OD1 application (100 nM) for 5 min. (**ii)** Group data showing that OD1 (100 nM) sensitized vaginal afferents to focal compression (*P<0.05, paired student t-test). **(C)** Representative traces showing the response to circular stretch (5 g) of vaginal afferent before and after OD1 (100 nM). Note that incubation with OD1 caused the afferent to continue to fire APs after cessation of the mechanical stimuli. (**ii-iii)** Grouped data showing OD1-induced sensitisation of vaginal afferents to circular stretch (5 g). OD1 induced increases in vaginal afferent firing to circular stretch were evident by (**ii)** an increased number of APs fired (*P<0.05, paired student t-test), and (**iii)** reduced time to elicit AP firing (*P<0.05, paired student t-test) generated after OD1. Grouped data are from n = 6 afferents from N = 3 Sham control mice. Data are mean ±SEM.

### Inhibition of Na_V_1.7 does not alter the mechanosensitivity of vaginal afferents, nor pain to vaginal distension in conscious Sham control mice

We next used a Tsp1a [41], a selective inhibitor of Na_V_1.7, to determine whether this channel contributes to vaginal afferent responsiveness to mechanical stimuli, *ex vivo*. In Sham control mice, exposure of vaginal afferent endings to Tsp1a [41] did not affect vaginal afferent responses to mucosal stroking (**Figure 2Ai-ii**), focal compression (**Figure 2Bi-ii**), and circular stretch (**Figure 2Ci-iii**).

**Figure 2:**
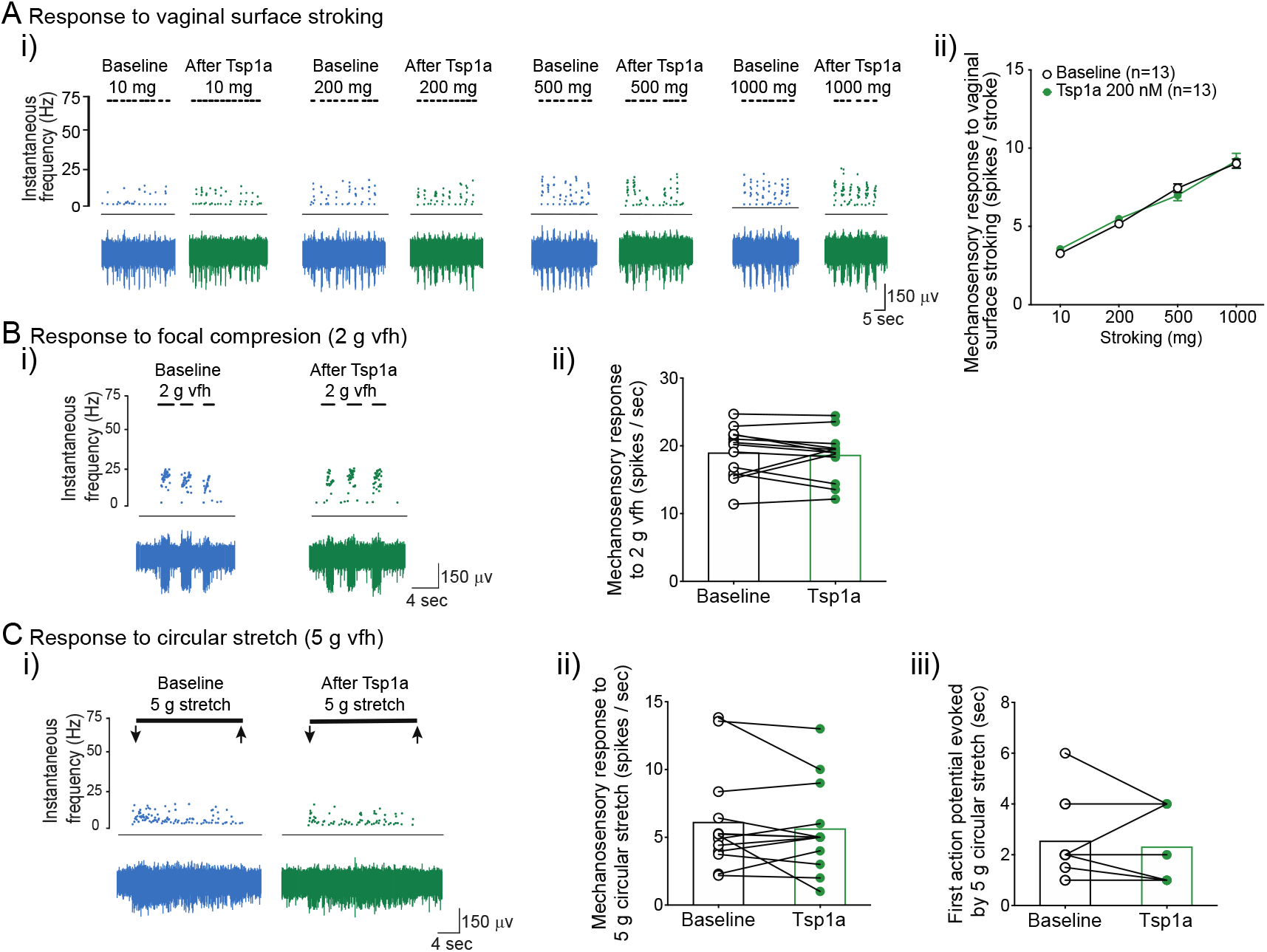
Inhibition of Na_V_1.7 with Tsp1a does not alter the responsiveness of pelvic vaginal afferents from Sham control mice to mechanical stimuli. (**A**) (i) Representative traces from ex vivo single-unit recordings of vaginal afferent from Sham control mice showing responses to vaginal stroking at baseline and after incubation with the Na_V_1.7 inhibitor Tsp1a (200 nM). (ii) Group data showing a lack of effect of Tsp1a (200 nM) on vaginal afferent sensitivity to receptive field stroking (P>0.05, two-way ANOVA followed by Bonferroni post-hoc comparison tests). (**B**) (i) Representative traces of vaginal afferent responses to focal compression with a 2 g vfh at baseline and in the presence of Tsp1a (200 nM). (ii) Group data showing a lack of effect of Tsp1a (200 nM) on vaginal afferent sensitivity to 2g vfh probing (P>0.05, paired student t-test). (**C**) (i) Representative traces of vaginal afferent responses to circular stretch (5 g) at baseline and in the presence of Tsp1a (200 nM). (ii) Group data showing a lack of effect of Tsp1a (200 nM) on vaginal afferent sensitivity to circular stretch of the vagina (P>0.05, paired student t-test). (iii) Additionally, Tsp1a did not change the time taken by vaginal afferents to fire their first AP in response to circular stretch (P>0.05, paired student t-test). Grouped data are from n = 13 afferents from N = 4 Sham control mice. Data are mean ±SEM.

We then investigated whether Tsp1a could modulate pain sensitivity evoked by vaginal distension (VD) in conscious Sham control mice. *In vivo* vaginal pain sensitivity was monitored by measuring the VMR to increasing VD pressures using EMG electrodes surgically implanted into the abdominal muscles as previously described [16; 17; 48]. In Sham control mice, we found that VD evoked an increase in the VMR, with the degree of VMR related to the amount of pressure applied to the vagina (**Figure 3A-B**). The *in vivo* sensitivity to pain evoked by VD in Sham control mice administered Tsp1a intravaginally was equal in magnitude to that displayed by mice treated with intravaginal saline (**Figure 3Ci**). This effect was observed across both non-noxious and noxious distension pressures (**Figure 3Cii-iii)**. Collectively, these results suggest that while activation of Na_V_1.7 can enhance afferent hypersensitivity in control states, inhibition of Na_V_1.7 does not alter the capacity of vaginal afferents to sense non-noxious and noxious stimuli.

**Figure 3:**
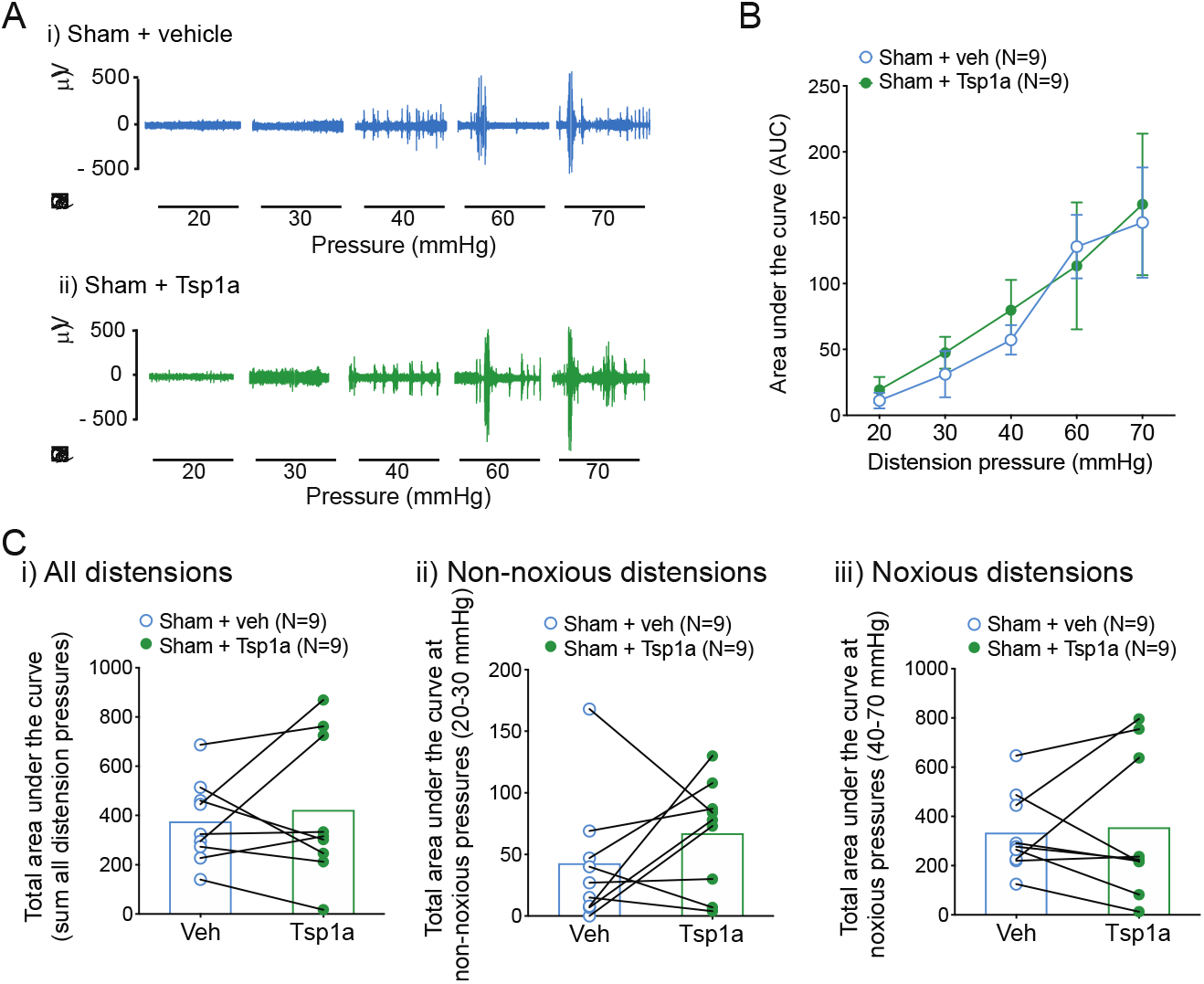
In vivo inhibition of Na_V_1.7 with Tsp1a does not alter pain evoked by vaginal distension in Sham control mice. **(A)** Representative EMG recordings at increasing VD pressures (horizontal bars represent 30 s VD at pressures of 20–70 mmHg) in conscious Sham control mice following intravaginal administration of (**i)** vehicle (saline) followed by (**ii)** Tsp1a (200 nM). **B)** In Sham control mice, no significant difference was observed in VMRs between intravaginal vehicle treatment or subsequent intravaginal treatment with Tsp1a (P > 0.05 generalised estimating equations followed by LSD post-hoc test). (**C)** Grouped data showing the lack of effect of Tsp1a evidenced by the total AUC of the VMRs (**i)** to all at all distensions pressures (sum of the AUCs obtained at each distension pressure), and the AUC of the VMRs at (**ii)** non-noxious distension pressures (20–30 mmHg, P > 0.05, paired student t-test), and (**iii)** noxious distension pressures (40–70 mmHg, P > 0.05, paired student t-test). Group data are from N = 9 Sham control mice that received vehicle treatment followed by Tsp1a treatment. Data are mean ±SEM.

### Vaginal afferents from mice with endometriosis display hypersensitivity to mechanical stimuli

We next investigated whether Na_V_1.7 plays a functional role in nociceptive signalling associated with endometriosis. We previously showed that, in a mouse model of IBS, the sensory afferents innervating the colon develop mechanical hypersensitivity. These hypersensitive colonic afferents send enhanced nociceptive signals to the CNS, ultimately causing increased pain sensitivity to colorectal distension *in vivo* [13; 15; 31; 33; 35]. Although similar neuroplasticity of sensory afferents has been suggested to occur in endometriosis [2; 8; 47], this has not been demonstrated experimentally. Therefore, we investigated whether sensory afferents innervating the vagina of mice with endometriosis become hypersensitive to mechanical stimuli.

Using our *ex vivo* vaginal afferent recordings, we characterised the response of vagina-innervating afferents to three different mechanical stimuli, in both Sham control mice and mice with fully developed endometriosis (Endo) **(Figure 4)**. Compared to Sham control mice, vaginal afferents from Endo mice fired significantly more APs in response to vaginal surface stroking and circular stretch (**Figure 4A and C**). Moreover, AP firing evoked by circular stretch commenced more rapidly within afferents from Endo mice (**Figure 4Civ**). Overall, these data indicate that the development of endometriosis enhances the sensitivity of pelvic vaginal afferents to mechanical stimuli.

**Figure 4:**
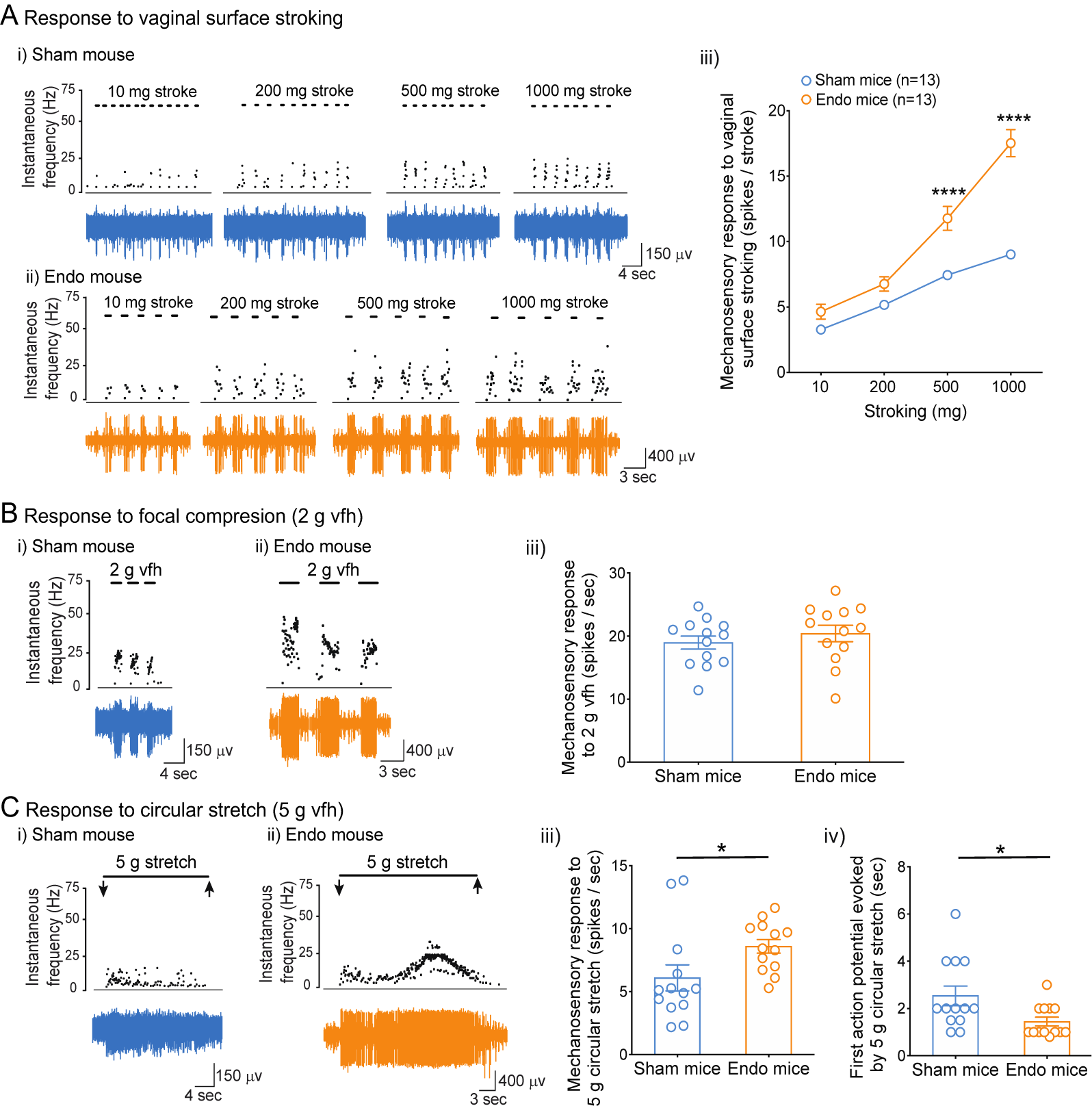
Pelvic vaginal afferents from endometriosis mice display hypersensitivity to mechanical stimuli. **(A)** Representative traces of vaginal afferents from (**i)** Sham control mice and (**ii)** endometriosis (Endo) mice showing baseline AP discharge in response to graded stroking of the vaginal surface. (**iii)** Group data showing that vaginal afferents from Endo mice display increased sensitivity to graded stroking of the vaginal surface (****P < 0.0001 at 500 mg and 1000 mg, two-way ANOVA followed by Bonferroni post-hoc comparison tests). (**B)** Representative traces of vaginal afferents from (**i)** Sham control mice and (**ii)** Endo mice showing baseline responses to focal compression with a 2 g vfh. (**iii)** Group data showing a lack of difference between groups in response to vaginal afferent focal compression (P > 0.05, two-tailed unpaired student t-test). (**C)** Representative traces of vaginal afferents from (**i)** Sham control mice and (**ii)** Endo mice showing baseline responses to circular stretch (5 g). Compared with Sham controls, vaginal afferents from Endo mice showed hypersensitivity to circular stretch, as determined by (**iii)** an increased number of APs generated by circular stretch (*P<0.05, two-tailed unpaired student t-test), and (**iv)** reduced latency required for AP firing (*P<0.05, two-tailed unpaired student t-test). Grouped data are from n = 13 afferents from N = 4 Sham control mice, and n = 13 afferents from N = 4 Endo mice. Data are mean ±SEM.

### Inhibition of Na_V_1.7 reduces endometriosis-associated vaginal afferent hypersensitivity

We then examined whether selective inhibition of Na_V_1.7 with Tsp1a [41] was able to reverse endometriosis-associated vaginal afferent hypersensitivity. We found that exposure of vaginal afferent endings to Tsp1a significantly reduced vaginal afferent firing to vagina surface stroking (**Figure 5Ai-iii**), focal compression (**Figure 5Bi-iii)**, and circular stretch (**Figure 5Ci-iii)** in Endo mice. Moreover, the time taken for these afferents to fire the first AP was significantly longer after incubation with Tsp1a (**Figure 5Civ)**. Overall, these data indicate that Na_V_1.7 activity contributes to the enhanced sensitivity developed in vaginal afferents from Endo mice.

**Figure 5:**
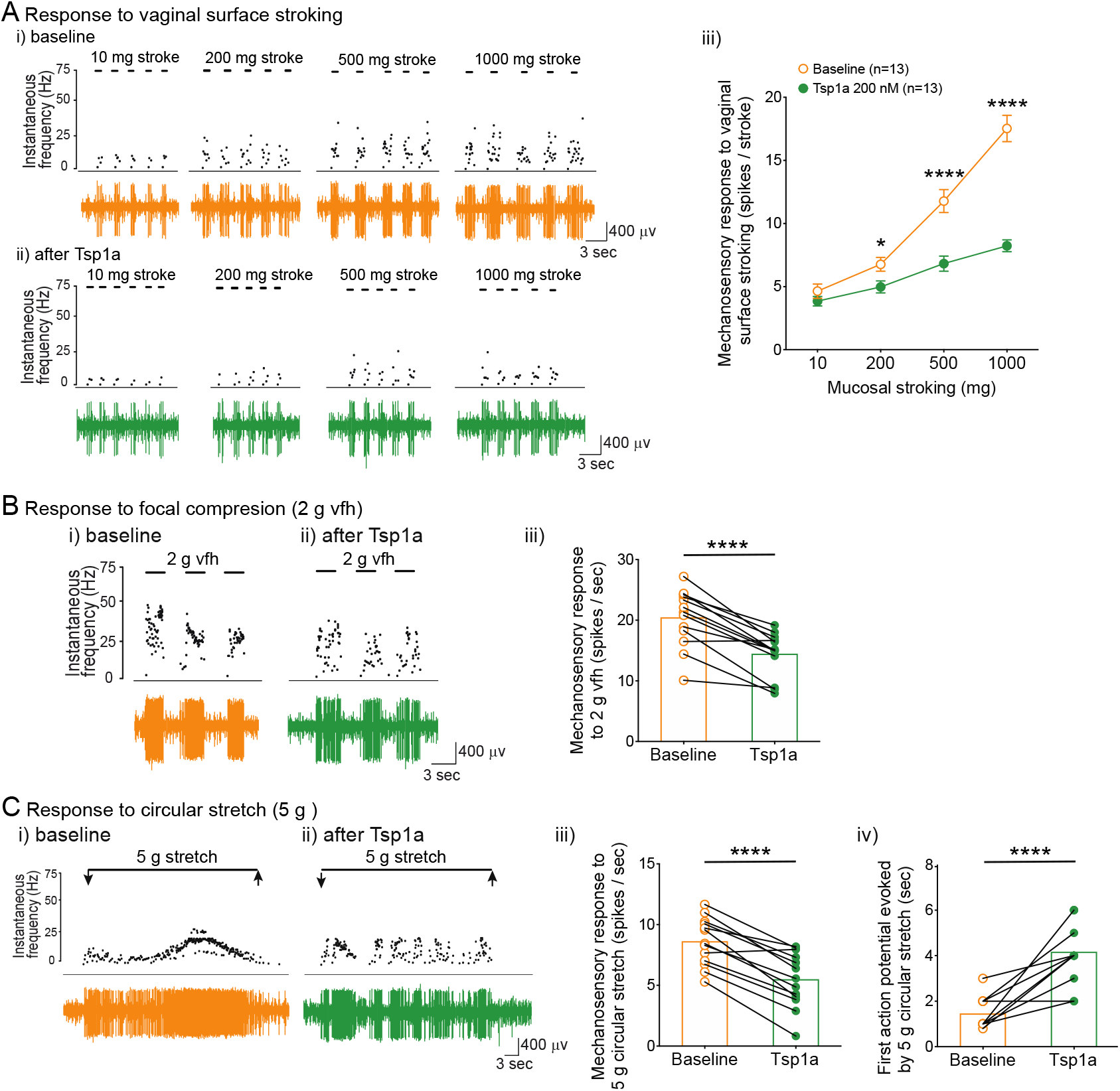
Vaginal afferents from endometriosis mice display reduced responsiveness to mechanical stimuli following inhibition of Na_V_1.7 with Tsp1a. **(A)** Representative traces of vaginal afferents from endometriosis mice showing responses to vaginal stroking at (**i)** baseline and (**ii)** after incubation with the Na_V_1.7 inhibitor Tsp1a (200 nM). (**iii)** Grouped data showing Tsp1a significantly reduces vaginal afferent sensitivity to stroking (*P<0.05 at 200 mg, and ****P<0.0001 at 500 mg and 1000 mg, two-way ANOVA followed by Bonferroni post-hoc comparison tests). (**B)** Representative traces of vaginal afferents from endometriosis mice showing responses to focal compression at (**i)** baseline and (**ii)** after incubation with the Na_V_1.7 inhibitor Tsp1a (200 nM). (**iii)** Group data showing Tsp1a significantly reduces vaginal afferent sensitivity to probing (****P<0.0001, paired student t-test. (**C)** Representative traces of vaginal afferents from endometriosis mice showing responses to circular stretch at (**i)** baseline and (**ii)** after incubation with the Na_V_1.7 inhibitor Tsp1a (200 nM). **iii)** Group data showing Tsp1a significantly reduces vaginal afferent sensitivity to circular stretch of the vagina (****P<0.0001, paired student t-test). (**iv)** Additionally, Tsp1a significantly increased the time taken by vaginal afferents to fire their first AP in response to circular stretch **(Civ)** (****P<0.0001, paired student t-test). Grouped data are from n = 13 afferents from N = 4 Sham control mice. Data are mean ±SEM.

### Inhibition of Na_V_1.7 reverses endometriosis-associated chronic pelvic pain to vaginal distension (VD)

We then investigated whether the observed alteration in sensory signalling from the vagina to the CNS translates to changes in pain sensitivity evoked by vaginal distension in conscious mice with endometriosis. We first confirmed our previous findings and demonstrated that Endo mice display enhanced pain sensitivity to VD [17]. Here we found that, compared to Sham control mice, Endo mice had elevated VMR responses to VD distension at all pressures tested (20–70 mmHg) (**Figure 6A-C**). Further analysis revealed the VMR to non-noxious (20–30 mmHg) (**Figure 6Cii)** and noxious (40– 70 mmHg) (**Figure 6Ciii)** DPs were significantly enhanced in Endo mice. Endo mice displayed both allodynia and hyperalgesia, which are hallmark signs of CPP in a wide range of inflammatory diseases, including endometriosis [6; 7; 17; 47; 48; 50; 58; 62]. We then found that Endo mice treated intravaginally with Tsp1a displayed significantly reduced VMRs evoked by VD compared with Endo mice treated intravaginally with vehicle (**Figure 7A-C**). This effect was observed at all non-noxious and noxious VD pressures that we tested (**Figure 7Cii-iii)**. Interestingly, Tsp1a reduced the VMRs in Endo mice to levels observed in Sham controls, suggesting that pharmacological inhibition of Na_V_1.7 on vaginal afferents is an effective strategy to reverse the vaginal allodynia and hyperalgesia associated with endometriosis.

**Figure 6.**
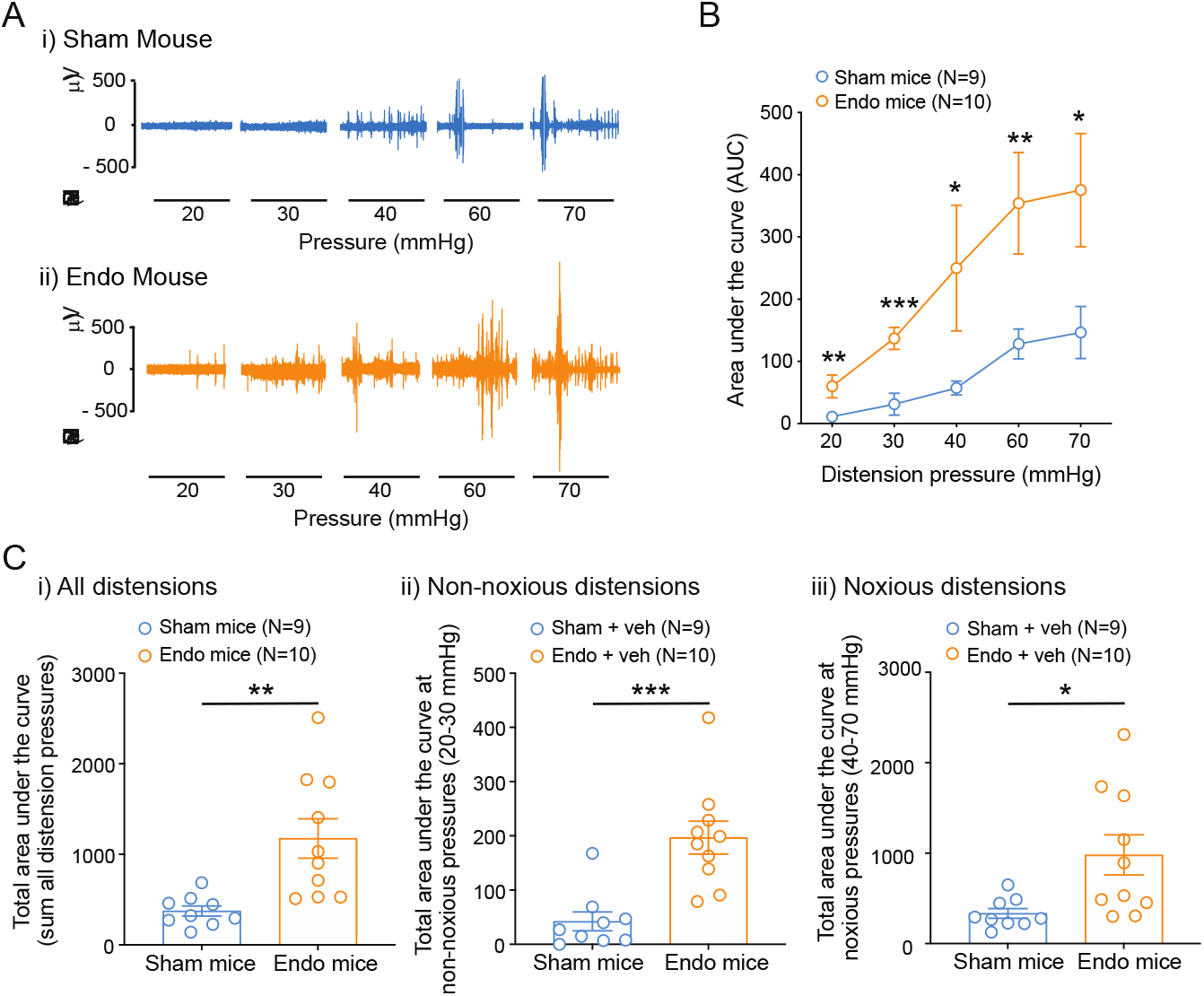
Mice with endometriosis display increased pain (allodynia and hyperalgesia) evoked by vaginal distension in vivo. **(A)** Representative EMG recordings showing the VMR to vaginal distension in (**i)** a Sham control mouse and (**ii)** a mouse with endometriosis. (**B)** Grouped AUC data showing that Endo mice displayed significantly en(hanced VMRs compared to their Sham counterparts (*P<0.05, **P<0.01, ***P<0.001, generalised estimating equations followed by LSD post-hoc test). (**C)** Grouped data showing that Endo mice have significantly enhanced total AUC compared with Sham control mice. Total AUC was obtained by combining the AUCs of the VMRs obtained at each distension pressure (**P<0.01, two-tailed unpaired student t-test). **D)** Further analysis reveals that mice with endometriosis have increased VMRs to both (**i)** non-noxious (20–30 mmHg, ***P<0.001, two-tailed unpaired student t-test), and (**ii)** noxious (40–70mmHg, *P<0.05, two-tailed unpaired student t-test) distension pressures. Grouped data are from N = 9 Sham control mice and N = 10 Endo mice. Data are mean ±SEM.

**Figure 7.**
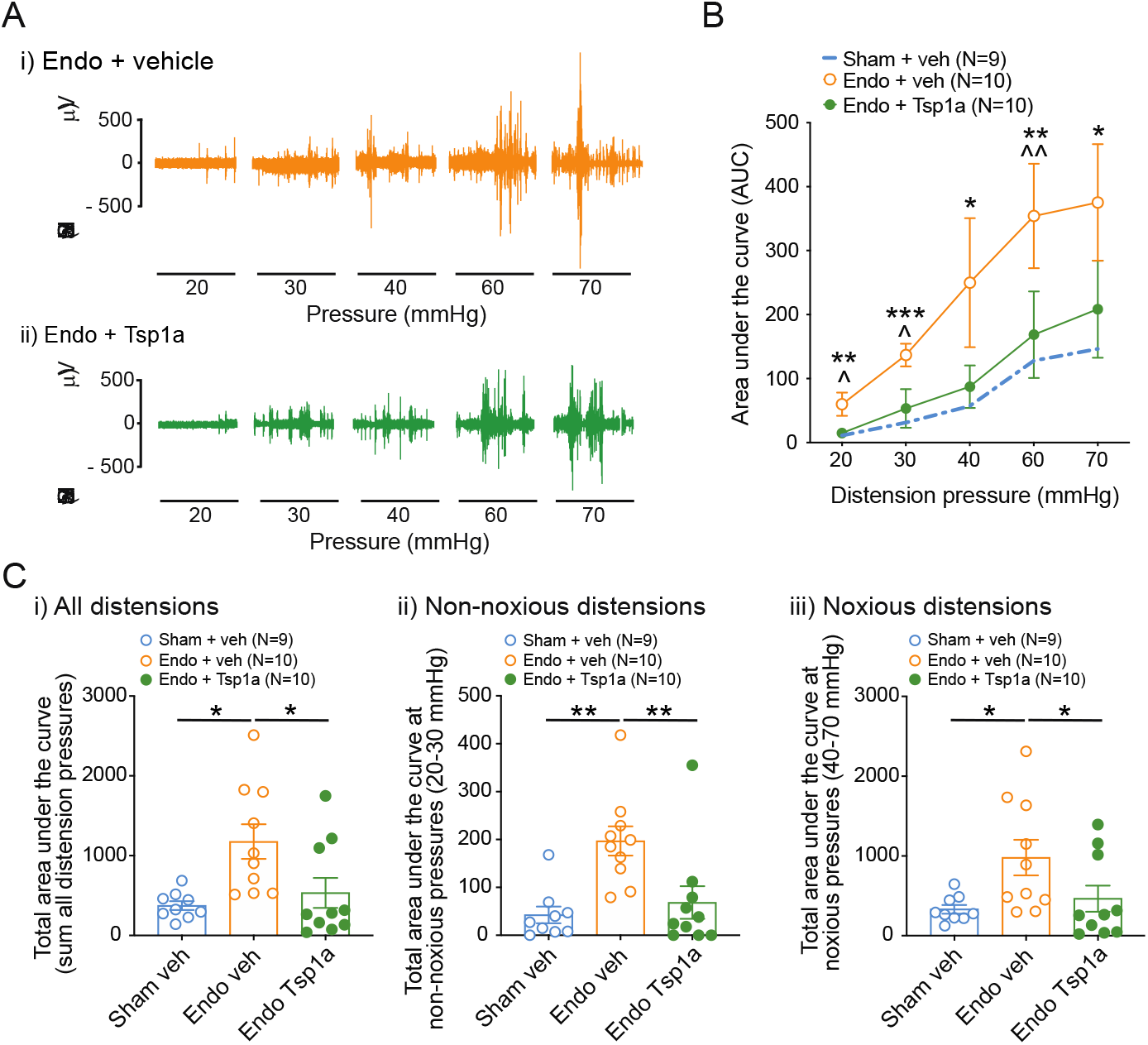
In endometriosis mice inhibition of Na_V_1.7 with Tsp1a reduces allodynia and hyperalgesia to vaginal distension. **(A)** Representative examples of EMG responses to VD in Endo mice following intravaginal treatment with (**i)** vehicle or (**ii)** Tsp1a. (**B)** Grouped data showing that the enhanced VMRs displayed by Endo mice (Endo + veh, orange) is significantly reduced by intravaginal treatment with 200 nM Tsp1a (Endo + Tsp1a, green). Note that the VMRs of Endo mice treated with Tsp1a are comparable to those displayed by Sham control mice (Sham + veh, blue dotted line). Statistical differences between Sham and Endo mice treated with vehicle are denoted by an asterisk (*P<0.05, **P<0.01, ***P<0.001, generalised estimating equations followed by LSD post hoc-test); whilst ^ indicates statistical differences between Endo mice treated with vehicle or Tsp1a (^P<0.05 and ^^P<0.01, generalised estimating equations followed by LSD post-hoc test). (**C)** Grouped data showing intra-vaginal treatment of Endo mice with Tsp1a significantly reduces (**i)** the elevated total AUC (sum of the AUCs obtained at each distension pressure), normalising these responses to those comparable of Sham control mice (*P<0.05, One-way ANOVA with Kruskai-wallis post-hoc test). Further analysis of the AUC data shows that Tsp1a was effective in reducing VMRs at both (**ii)** non-noxious (20–30 mmHg, **P<0.01, One-way ANOVA with Kruskal-Wallis post-hoc test); and (**iii)** noxious distension pressures (40–70 mmHg, *P<0.05, One-way ANOVA with Kruskal-Wallis post-hoc test). Grouped data are from N = 9 Sham control mice treated with vehicle, N = 10 Endo mice treated with vehicle and Tsp1a. Data are mean ±SEM.

## Discussion

Endometriosis is a chronic disorder characterised by infertility and CPP. Currently there is a lack of effective analgesic treatments for endometriosis-associated CPP mainly due to our lack of knowledge about its aetiology and pathogenesis [43; 61]. How sensory neurons innervating the female reproductive tract detect and transmit pain and how they are altered in endometriosis remains poorly understood. We recently showed that all nine members of the Na_V_ channel family are expressed in pelvic vaginal afferents, although their relative expression in these neurons varies greatly. The pan-Na_V_ channel agonist, veratridine, can drive vaginal afferent hypersensitivity, whilst the pan-Na_V_ channel blocker TTX reduces vaginal afferent sensitivity. These findings demonstrate that Na_**V**_ channels play an important role in regulating vaginal sensation in healthy animals and point towards a key role of TTX-sensitive Na_**V**_ channels [16]. However, the specific identity of the TTX-sensitive Na_V_ channels contributing to vaginal nociceptive signalling and their contribution to endometriosis-associated CPP has remained unclear.

In this study we addressed this by investigating the role of Na_V_1.7 in normal vaginal mechanosensitivity and endometriosis-associated vaginal mechanical hypersensitivity. We decided to target this channel because: (i) Na_V_1.7 is TTX-sensitive; (ii) it is abundantly expressed (98–100%) in sensory neurons that innervate pelvic organs affected by endometriosis, including the vagina [16], colon [38], and bladder [32]; (iii) we have unique access to selective modulators of Na_V_1.7 [23; 41]; (iv) we recently demonstrated a role for Na_V_1.7 in chronic visceral pain [41]; (v) various Na_V_1.7-selective inhibitors are in clinical trials for various types of pain [1; 3; 29; 44; 60].

First, we demonstrated that activation of Na_V_1.7 with OD1, a α-highly selective Na_V_1.7 agonist [23], dramatically enhanced the responsiveness of vaginal afferents from Sham control mice to three different types of mechanical stimuli. Interestingly, we observed that whilst OD1 failed to elicit AP discharge in the absence of mechanical stimuli (direct activation of afferents), half of the afferents incubated with OD1 continued to fire APs after cessation of the mechanical stimulus. This observation could be explained by the mechanism by which OD1 alters Na_V_1.7 channel function. Kinetically, Na_V_1.7 is slow to close and subsequently inactivates following its activation [34; 42]. OD1 inhibits the fast inactivation of Na_V_1.7, which ultimately disrupts the resting threshold equilibrium, maintaining hyperexcitable sensory neurons in the absence of further stimuli [23; 42; 49]. This is particularly relevant in genetic gain-of-function mutations, where the enhanced role of Na_V_1.7 in sensory neurons maintains them in a hyperexcitable state, increasing AP firing and ultimately driving sensory signalling in the absence of stimuli [22; 42]. Overall, our findings indicate that vaginal afferent responses to mechanical stimuli in control conditions can be enhanced by pharmacological activation of Na_V_1.7.

We next examined whether selective inhibition of Na_V_1.7 could reduce the ability of vaginal afferents to sense baseline mechanical stimuli in Sham control mice. For this we used the peptide Tsp1a, a selective inhibitor of Nav1.7 derived from venom of the Peruvian tarantula *Thrixopelma spec*. [41]. We found that Tsp1a does not alter the ability of vaginal afferents to respond to mechanical stimuli *ex vivo*. These findings were further supported by *in vivo* studies, whereby intravaginal application of Tsp1a failed to alter vaginal sensitivity to distension in conscious Sham control mice. Overall, our results are consistent with previous reports indicating that Na_V_1.7 does not appear to contribute to baseline visceral nociception in healthy conditions [36; 41].

However, a possible role for Na_V_1.7 in endometriosis-associated CPP has previously been suggested. There is evidence of *SCN9A* (the gene encoding Na_V_1.7) overexpression in endometriosis lesions collected from women with pain, compared to women without endometriosis/without pain [30]. Notably, human embryonic stem cells with a nociceptive phenotype treated with the estrogen receptor β agonist 2,3-bis(4-hydroxy-phenyl)-propionitrile display enhanced expression of *SCN9A* [30]. Furthermore, macrophages activated with peritoneal fluid from women with endometriosis can also enhance the expression of *SCN9A* in cultured human sensory neurons *in vitro*, via a macrophage-derived insulin-like growth factor 1 (IGF-1) mechanism [27]. While changes in the expression profile of Na_V_1.7 have been associated with endometriosis in these studies, the functional role of Na_V_1.7 in sensing pain from pelvic organs affected by endometriosis has not yet been explored. Therefore, we investigated the role of Na_V_1.7 in our established syngeneic mouse model of endometriosis [48]. We show here, for the first time, that pelvic afferents innervating the mouse vagina become hypersensitive to mechanical stimuli in endometriosis. This hypersensitivity was also displayed *in vivo*, with conscious Endo mice displaying both allodynia and hyperalgesia to vaginal distension compared to their Sham control counterparts. These findings demonstrate that increased nociceptive signalling from the vagina results in increased pain sensitivity *in vivo*, which closely corresponds with nociceptive mechanisms observed in other visceral organs [13; 15; 31; 33; 35].

Importantly, inhibition of Na_V_1.7 with Tsp1a was effective in reducing *ex vivo* vaginal afferent hypersensitivity in mice with endometriosis. This attenuation was also apparent *in vivo*; the VMR evoked by vaginal distension in conscious Endo mice was reversed with a single intravaginal dose of Tsp1a, with sensitivity restored to the levels seen in Sham control animals. Notably, this effect was apparent at non-noxious and noxious distension pressures, suggesting that Tsp1a is able alleviate vaginal allodynia and hyperalgesia in mice with endometriosis. Overall, our findings are consistent with the notion that enhanced activity of Na_V_1.7 is able to drive, and contribute to, heightened pain and nociception in chronic disease states [34; 41; 57]. However, Na_V_1.7 does not appear to contribute to visceral nociception in healthy conditions [36; 41]. This is an important concept, because an ideal analgesic should target the key mechanisms responsible for the pathological pain underlying the condition and provide pain relief without the loss of baseline sensory functions, which are essential for host responses to the external and internal environments.

Although Na_V_1.7 is a widely accepted as pain target, the translation to clinical significance as a treatment for chronic pain has been lacking [24]. This is likely due to a multiplicity of factors including: (i) genetic mutations impacting pain sensitivity in humans; (ii) the lack of selectivity and tissue penetration of some Na_V_1.7 inhibitors, and (iii) the underestimated impact of the effect of the auxiliary β subunits on Na_V_1.7 pharmacology [24]. Our findings showing different roles for Na_V_1.7 in nociceptive signalling in health and endometriosis states are in keeping with recent studies. For example, inhibition of Na_V_1.7 does not alter colonic afferent signalling in healthy states [36; 41], whereas Na_V_1.7 inhibition in a mouse model of IBS reverses mechanical hypersensitivity *ex vivo* and *in vivo* [41]. Other studies show that indirect targeting of Na_V_1.7 activity in sensory neurons by specifically inhibiting Na_V_1.7 trafficking and surface expression with compound 194, is effective at reversing thermal and mechanical hypersensitivity in a rodent model of neuropathic pain [28; 46]. Overall, the role of Na_V_1.7 in pain is complex, with somatosensory studies suggesting that Na_V_1.7 blockers alone may not replicate the analgesic phenotype of Na_V_1.7^-/-^ mutant mice and humans with congenital insensitivity to pain due to loss-of-function mutations in *SCN9A*. However, their effects may be potentiated with exogenous opioids [52] and G-protein coupled receptors (GPCRs) [39; 53]. For example, some studies have shown that Na_V_1.7 regulates the efficacy and balance of GPCR-mediated pro- and anti-nociceptive intracellular signalling, so that without Na_V_1.7 the balance of these processes is shifted toward anti-nociception [39]. Other animal studies show that post-surgical pain can be successfully treated with Na_V_1.7 inhibitors alone or at subtherapeutic doses in combination with baclofen or opioids, suggesting super-additive antinociceptive effects when Na_V_1.7 inhibitors are applied in combination with baclofen or opioids [53]. Whether opioidergic and GPCR mechanisms contribute to Na_V_1.7’s role in endometriosis-associated CPP will be the subject of future studies. However, it is interesting to note that several GPCR and opioidergic mechanisms are upregulated in colonic nociceptors [10-12; 14; 21; 37; 54; 55] in IBS models and that inhibition of Na_V_1.7 also reverses pathological pain in these IBS models [41].

In conclusion, our study demonstrates that Na_V_1.7 contributes to the vaginal hyperalgesia associated with endometriosis. Moreover, it reveals that pharmacological inhibition Na_V_1.7 with Tsp1a could be a viable therapeutic option to treat endometriosis-associated CPP. Together with our recently published findings showing that Tsp1a could reverse colonic hypersensitivity in a mouse model of IBS [41], these results suggest that pharmacological targeting of Na_V_1.7 is a valid approach to treat CPP associated with visceral pain disorders including endometriosis.

## Author contributions

J.C, J.M, and S.M.B contributed to experimental design, analysis, and intellectual input. J.C and J.M performed the experiments described in this paper. I.V and G.F.K provided peptides targeting Na_V_ channels and intellectual input. J.C made the figures and wrote the manuscript with contributions from all authors.

## Acknowledgments

We acknowledge funding from the National Health and Medical Research Council of Australia (Ideas Grant APP1181448 to J.C.; Investigator Leadership Grant APP2008727 to S.M.B.; Principal Research Fellowship APP1136889 to G.F.K.; Career Development Fellowship APP1162503 to I.V; Development Grant APP2014250 to S.M.B. and G.F.K.), the Australian Research Council (Discovery Project DP220101269 to S.M.B.; Centre of Excellence grant CE200100012 to G.F.K.) and The Hospital Research Foundation (PhD Scholarship SAPhD000242018 to J.M.). None of the authors have declared conflict of interest.

## Methods

### 2.1. Animals

All experiments involving animals were approved by the Animal Ethics Committee of the South Australian Health and Medical Research Institute (SAHMRI; ethics number SAM342) and conformed to the relevant regulatory standards and ARRIVE guidelines. Female C57BL/6J mice at 8–13 weeks of age were used and acquired from an in-house C57BL/6J breeding program (JAX strain #000664; originally purchased from The Jackson Laboratory (breeding barn MP14; Bar Harbor, ME; USA)) within SAHMRI’s specific and opportunistic pathogen-free animal care facility. Mice were group-housed (maximum five mice per cage) within individual ventilated cages (IVC), which were filled with coarse chip dust-free aspen bedding (PuraChips Aspen coarse 63L; Cat# ASPJMAEB-CA, Able Scientific, Australia). These cages were stored on ventilated IVC racks in specific housing rooms within a temperature-controlled environment of 22°C and a 12 h light/12 h dark cycle. Mice had free access to LabDiet® JL Rat and Mouse/Auto6F chow (Speciality Feeds, Australia) and autoclaved reverse osmosis purified water. Female mice were group housed in IVC cages and the littermate male mice were separated at weaning. All female mice use in this study were unmated virgins.

### 2.2. Pharmacological modulators

The α-scorpion toxin OD1, a selective agonist of Na_V_1.7, was produced via solid-phase peptide synthesis (SPPS) as previously described [23]. The spider-venom peptide Tsp1a, a selective inhibitor of Na_V_1.7, was also produced via SPPS as previously described [41].

### 2.3. Syngeneic inoculation mouse model of endometriosis

In this study we used a syngeneic mouse model of endometriosis, previously established and characterised by our group [48]. Briefly, female mice were ovariectomised under isoflurane and received estrogen (100 μg/kg estradiol benzoate) intraperitoneally (IP). Following ovariectomy recovery (minimum of 5 days), endometriosis (Endo) or Sham inductions were performed as previously described [48]. For this, recipient mice were anesthetized under isoflurane, and a small (0.5 cm) incision was made into the peritoneal space just below the umbilicus. 250 μL of minced endometrial tissue suspension obtained from donor mice (in PBS plus penicillin: 100 U/ml and streptomycin: 100 μg/ml) was then inoculated into the peritoneal cavity using a 1-ml pipette, as described previously [48]. A ratio of 1 donor mouse per 2 recipient mice was used. The incision was then closed using 6.0 Prolene sutures, and a gentle massaging of the abdominal cavity was performed to help disperse the inoculated endometrial fragments. Sham surgeries were performed by inoculating 250 μL of sterile PBS plus penicillin (100 U/ml) and streptomycin (100 μg/ml) in the absence of any tissue. All recipient mice received a low dose (0.05 mg/kg) of analgesic buprenorphine prior to the commencement of surgery. Throughout the surgery and during the recovery period, animals were kept on a heating pad to maintain body temperature and monitored daily (for 5 consecutive days) for postsurgical complications. To maintain steady levels of circulating estrogen and minimize any difference related to the stage of the oestrous cycle, both Endo mice and Sham control mice were given IP injections of estrogen (100 μg/kg estradiol benzoate) immediately after surgery. All mice continued to receive estrogen IP once a week until full development of endometriosis (up to 8–10 weeks).

### 2.4 Ex vivo afferent recording from pelvic nerves innervating the female reproductive tract

Single-unit afferents recordings from the pelvic nerve innervating the vagina of Sham control mice and mice with endometriosis were performed using an *ex vivo* afferent recording preparation as previously described [16]. Briefly, the intact female reproductive tract was removed along with the attached neurovascular bundle containing the pelvic nerve and transferred to ice-cold Krebs solution (in mM: 117.9 NaCl, 4.7 KCl, 25 NaHCO_3_, 1.3 NaH_2_PO_4_, 1.2 MgSO_4_∙(H_2_O)7, 2.5 CaCl_2_, 11.1 D-glucose). Tissue was opened longitudinally and pinned flat, mucosal side up, in a specialised organ bath consisting of two adjacent compartments generated from clear acrylic (Danz Instrument Service, Adelaide, South Australia, Australia), the floors of which were lined with Sylgard (Dow Corning Corp., Midland, MI). The neurovascular bundle containing the pelvic nerve was extended from the tissue compartment into the paraffin oil filled recording compartment where they were laid onto a mirror. The organ compartment was perfused with Krebs solution and bubbled with carbogen (95% O_2_, 5% CO_2_) at a temperature of 34°C. The pelvic nerve was dissected away from the neurovascular tissue and divided into 6–10 bundles. One of the bundles was placed onto a platinum recording electrode. A separate platinum reference electrode rested on the mirror in a small pool of Krebs solution adjacent to the recording electrode. Action potentials, generated by mechanical stimuli applied to the afferent’s receptive field, were recorded using a differential amplifier, and filtered and sampled (20 kHz) using a 1401 interface (Cambridge Electronic Design, Cambridge, UK).

#### 2.4.1. Mechanosensory profile of pelvic afferents innervating the vagina

Receptive fields tested in this study were limited to the vagina (above vaginal opening and below cervix) and were identified by systematically stroking the mucosal surface of the vagina with a stiff brush to activate all subtypes of vaginal mechanoreceptors. Mechanosensory properties of the pelvic afferents innervating a particular receptive field within the vagina were assessed by three distinct mechanical stimuli as previously described [16]. These included: (i) static probing with calibrated von Frey hairs (vfh; 2 g force; applied 3 times for a period of 3 s); (ii) mucosal stroking of the vaginal surface with calibrated vfh (10–1000 mg force; applied 10 times each); and (iii) circular stretch (5 g; applied for a period of 30 s). Once baseline mechanosensitivity was tested, a small chamber was then placed onto the mucosal surface of the vagina, surrounding the afferent receptive field. Residual Krebs solution within the chamber was aspirated and the Na_V_ channel modulators OD1 (100 nM) and Tsp1a (200 nM) were applied in separate experimental preparations for 5 min each. The afferent receptive field was then re-tested using the same three mechanical stimuli.

#### 2.4.2. Statistical analysis of afferent recording data

Action potentials were analysed off-line using Spike 2 (version 5.21) software (Cambridge Electronic Design, Cambridge, UK) and discriminated as single units based on distinguishable waveforms, amplitudes, and durations. Data are expressed as mean ± SEM. n = the number of afferents recorded, N = the number of animals used for those specific experiments. Data were statistically compared using Prism 7 software (GraphPad Software, San Diego, CA, USA) and, where appropriate, analysed using paired or un-paired Student’s *t*-test and one- or two-way analysis of variance (ANOVA) with Bonferroni post hoc tests. Differences were considered statistically significant at *P* < 0.05.

#### 2.5 Assessment of vaginal pain sensitivity in vivo

We quantified the visceromotor response (VMR) to vaginal distension (VD) as an objective measurement of vaginal sensitivity to pain in fully awake mice as previously described [16; 17; 48].

##### 2.5.1. Surgical implantation of electromyography electrodes and VMR assessment

Electromyography (EMG) electrodes were implanted in the abdominal musculature. All mice received prophylactic antibiotics (Baytril® 5 mg/kg s.c.) and pain relief (buprenorphine 0.05 mg/kg s.c.). After surgery, animals were single housed to protect the EMG electrodes. Animals were allowed to recover from surgery for at least 3 days before VMR assessment. On the day of VMR assessment, animals were briefly sedated with isoflurane and Tsp1a (200 nM) or vehicle (saline) were administered intravaginally via a small canula inserted into the vaginal canal. Immediately after, a lubricated 3 mm length latex balloon was gently passed through the vagina and inserted up to 1 mm proximal to the vaginal verge and secured to the base of the tail. The balloon was then connected to a barostat (Isobar 3 Barostat, G&J Electronics) for pressure-controlled rapid inflation of air. Animals were transferred to a restrainer with dorsal access and the EMG electrodes were relayed to a data acquisition system. Vaginal distensions were applied by the barostat, ranging from the non-noxious to the noxious range (20-30-40-60-70 mmHg, 30 s duration, 3 min interval between distensions), after animals regained consciousness (∼ 10 min drug administration and balloon insertion). The corresponding EMG signal was recorded (NL100AK headstage), amplified (NL104), filtered (NL 125/126, Neurolog, Digitimer Ltd, bandpass 50–5000 Hz), and digitised (CED 1401, Cambridge Electronic Design, Cambridge, UK) to a PC for off-line analysis using Spike2 (Cambridge Electronic Design).

##### 2.5.2. Statistical analysis of VMR data

To quantify the magnitude of the VMR at each distension pressure, the area under the curve (AUC) during the distension (30 s) was corrected for the baseline activity (AUC pre-distension, 30 s). Total area under the curve was quantified by adding the individual AUC at each distension pressure. VMR data are presented as mean ± SEM. N represents the number of animals. VMR data were statistically analysed by generalised estimating equations followed by Fisher’s least significant difference (LSD) post-hoc test when appropriate using SPSS 23.0. Analysis and figures were prepared using Prism 7 (GraphPad Software, San Diego, CA, USA). Differences were considered statistically significant at P < 0.05.

## References

[1] Alexandrou AJ, Brown AR, Chapman ML, Estacion M, Turner J, Mis MA, Wilbrey A, Payne EC, Gutteridge A, Cox PJ, Doyle R, Printzenhoff D, Lin Z, Marron BE, West C, Swain NA, Storer RI, Stupple PA, Castle NA, Hounshell JA, Rivara M, Randall A, Dib-Hajj SD, Krafte D, Waxman SG, Patel MK, Butt RP, Stevens EB. Subtype-Selective Small Molecule Inhibitors Reveal a Fundamental Role for Nav1.7 in Nociceptor Electrogenesis, Axonal Conduction and Presynaptic Release. PLoS One 2016;11(4):e0152405.

[2] Arnold J, Barcena de Arellano ML, Ruster C, Vercellino GF, Chiantera V, Schneider A, Mechsner S. Imbalance between sympathetic and sensory innervation in peritoneal endometriosis. Brain Behav Immun 2012;26(1):132–141.

[3] Bagal SK, Chapman ML, Marron BE, Prime R, Storer RI, Swain NA. Recent progress in sodium channel modulators for pain. Bioorg Med Chem Lett 2014;24(16):3690–3699.

[4] Becker CM, Missmer SA, Zondervan KT. Endometriosis. Reply. N Engl J Med 2020;383(2):194.

[5] Bennett DL, Woods CG. Painful and painless channelopathies. Lancet Neurol 2014;13(6):587–599.

[6] Berkley KJ, Cason A, Jacobs H, Bradshaw H, Wood E. Vaginal hyperalgesia in a rat model of endometriosis. Neurosci Lett 2001;306(3):185–188.

[7] Berkley KJ, McAllister SL, Accius BE, Winnard KP. Endometriosis-induced vaginal hyperalgesia in the rat: effect of estropause, ovariectomy, and estradiol replacement. Pain 2007;132 Suppl 1:S150–S159.

[8] Berkley KJ, Rapkin AJ, Papka RE. The pains of endometriosis. Science 2005;308(5728):1587–1589.

[9] Broad LM, Mogg AJ, Beattie RE, Ogden AM, Blanco MJ, Bleakman D. TRP channels as emerging targets for pain therapeutics. Expert Opin Ther Targets 2009;13(1):69–81.

[10] Brust A, Croker DE, Colless B, Ragnarsson L, Andersson Å, Jain K, Garcia-Caraballo S, Castro J, Brierley SM, Alewood PF, Lewis RJ. Conopeptide-Derived κ-Opioid Agonists (Conorphins): Potent, Selective, and Metabolic Stable Dynorphin A Mimetics with Antinociceptive Properties. J Med Chem 2016;59(6):2381–2395.

[11] Castro J, Garcia-Caraballo S, Maddern J, Schober G, Lumsden A, Harrington A, Schmiel S, Lindstrom B, Adams J, Brierley SM. Olorinab (APD371), a peripherally acting, highly selective, full agonist of the cannabinoid receptor 2, reduces colitis-induced acute and chronic visceral hypersensitivity in rodents. Pain 2022;163(1):e72–e86.

[12] Castro J, Grundy L, Deiteren A, Harrington AM, O’Donnell T, Maddern J, Moore J, Garcia-Caraballo S, Rychkov GY, Yu R, Kaas Q, Craik DJ, Adams DJ, Brierley SM. Cyclic analogues of α-conotoxin Vc1.1 inhibit colonic nociceptors and provide analgesia in a mouse model of chronic abdominal pain. Br J Pharmacol 2018;175(12):2384–2398.

[13] Castro J, Harrington AM, Garcia-Caraballo S, Maddern J, Grundy L, Zhang J, Page G, Miller PE, Craik DJ, Adams DJ, Brierley SM. alpha-Conotoxin Vc1.1 inhibits human dorsal root ganglion neuroexcitability and mouse colonic nociception via GABAB receptors. Gut 2017;66(6):1083–1094.

[14] Castro J, Harrington AM, Garcia-Caraballo S, Maddern J, Grundy L, Zhang J, Page G, Miller PE, Craik DJ, Adams DJ, Brierley SM. α-Conotoxin Vc1.1 inhibits human dorsal root ganglion neuroexcitability and mouse colonic nociception via GABA(B) receptors. Gut 2017;66(6):1083–1094.

[15] Castro J, Harrington AM, Hughes PA, Martin CM, Ge P, Shea CM, Jin H, Jacobson S, Hannig G, Mann E, Cohen MB, MacDougall JE, Lavins BJ, Kurtz CB, Silos-Santiago I, Johnston JM, Currie MG, Blackshaw LA, Brierley SM. Linaclotide inhibits colonic nociceptors and relieves abdominal pain via guanylate cyclase-C and extracellular cyclic guanosine 3’,5’-monophosphate. Gastroenterology 2013;145(6):1334-1346 e1331-1311.

[16] Castro J, Maddern J, Erickson A, Caldwell A, Grundy L, Harrington AM, Brierley SM. Pharmacological modulation of voltage-gated sodium (NaV) channels alters nociception arising from the female reproductive tract. Pain 2021;162(1):227–242.

[17] Castro J, Maddern J, Grundy L, Manavis J, Harrington AM, Schober G, Brierley SM. A mouse model of endometriosis that displays vaginal, colon, cutaneous, and bladder sensory comorbidities. FASEB J 2021;35(4):e21430.

[18] Catterall WA. Voltage-gated sodium channels at 60: structure, function and pathophysiology. J Physiol 2012;590(11):2577–2589.

[19] Catterall WA, Goldin AL, Waxman SG. International Union of Pharmacology. XLVII. Nomenclature and structure-function relationships of voltage-gated sodium channels. Pharmacol Rev 2005;57(4):397–409.

[20] Chaban VV. Visceral sensory neurons that innervate both uterus and colon express nociceptive TRPv1 and P2X3 receptors in rats. Ethn Dis 2008;18(2 Suppl 2):S2-20-24.

[21] de Araujo AD, Mobli M, Castro J, Harrington AM, Vetter I, Dekan Z, Muttenthaler M, Wan J, Lewis RJ, King GF, Brierley SM, Alewood PF. Selenoether oxytocin analogues have analgesic properties in a mouse model of chronic abdominal pain. Nat Commun 2014;5:3165.

[22] Dib-Hajj SD, Rush AM, Cummins TR, Hisama FM, Novella S, Tyrrell L, Marshall L, Waxman SG. Gain-of-function mutation in Nav1.7 in familial erythromelalgia induces bursting of sensory neurons. Brain 2005;128(Pt 8):1847–1854.

[23] Durek T, Vetter I, Wang CI, Motin L, Knapp O, Adams DJ, Lewis RJ, Alewood PF. Chemical engineering and structural and pharmacological characterization of the alpha-scorpion toxin OD1. ACS Chem Biol 2013;8(6):1215–1222.

[24] Eagles DA, Chow CY, King GF. Fifteen years of NaV 1.7 channels as an analgesic target: Why has excellent in vitro pharmacology not translated into in vivo analgesic efficacy? Br J Pharmacol 2022;179(14):3592–3611.

[25] Eisenberg VH, Weil C, Chodick G, Shalev V. Epidemiology of endometriosis: a large population-based database study from a healthcare provider with 2 million members. BJOG 2018;125(1):55–62.

[26] Erickson A, Deiteren A, Harrington AM, Garcia-Caraballo S, Castro J, Caldwell A, Grundy L, Brierley SM. Voltage-gated sodium channels: (NaV)igating the field to determine their contribution to visceral nociception. J Physiol 2018;596(5):785–807.

[27] Forster R, Sarginson A, Velichkova A, Hogg C, Dorning A, Horne AW, Saunders PTK, Greaves E. Macrophage-derived insulin-like growth factor-1 is a key neurotrophic and nerve-sensitizing factor in pain associated with endometriosis. FASEB J 2019;33(10):11210–11222.

[28] Francois-Moutal L, Dustrude ET, Wang Y, Brustovetsky T, Dorame A, Ju W, Moutal A, Perez-Miller S, Brustovetsky N, Gokhale V, Khanna M, Khanna R. Inhibition of the Ubc9 E2 SUMO-conjugating enzyme-CRMP2 interaction decreases NaV1.7 currents and reverses experimental neuropathic pain. Pain 2018;159(10):2115–2127.

[29] Goldberg YP, Price N, Namdari R, Cohen CJ, Lamers MH, Winters C, Price J, Young CE, Verschoof H, Sherrington R, Pimstone SN, Hayden MR. Treatment of Na(v)1.7-mediated pain in inherited erythromelalgia using a novel sodium channel blocker. Pain 2012;153(1):80–85.

[30] Greaves E, Grieve K, Horne AW, Saunders PT. Elevated peritoneal expression and estrogen regulation of nociceptive ion channels in endometriosis. J Clin Endocrinol Metab 2014;99(9):E1738–1743.

[31] Grundy L, Erickson A, Brierley SM. Visceral Pain. Annu Rev Physiol 2019;81:261–284.

[32] Grundy L, Erickson A, Caldwell A, Garcia-Caraballo S, Rychkov G, Harrington A, Brierley SM. Tetrodotoxin-sensitive voltage-gated sodium channels regulate bladder afferent responses to distension. Pain 2018;159(12):2573–2584.

[33] Grundy L, Harrington AM, Castro J, Garcia-Caraballo S, Deiteren A, Maddern J, Rychkov GY, Ge P, Peters S, Feil R, Miller P, Ghetti A, Hannig G, Kurtz CB, Silos-Santiago I, Brierley SM. Chronic linaclotide treatment reduces colitis-induced neuroplasticity and reverses persistent bladder dysfunction. JCI Insight 2018;3(19).

[34] Hameed S. Nav1.7 and Nav1.8: Role in the pathophysiology of pain. Mol Pain 2019;15:1744806919858801.

[35] Harrington AM, Brierley SM, Isaacs N, Hughes PA, Castro J, Blackshaw LA. Sprouting of colonic afferent central terminals and increased spinal mitogen-activated protein kinase expression in a mouse model of chronic visceral hypersensitivity. J Comp Neurol 2012;520(10):2241–2255.

[36] Hockley JR, Gonzalez-Cano R, McMurray S, Tejada-Giraldez MA, McGuire C, Torres A, Wilbrey AL, Cibert-Goton V, Nieto FR, Pitcher T, Knowles CH, Baeyens JM, Wood JN, Winchester WJ, Bulmer DC, Cendan CM, McMurray G. Visceral and somatic pain modalities reveal NaV 1.7-independent visceral nociceptive pathways. J Physiol 2017;595(8):2661–2679.

[37] Hughes PA, Castro J, Harrington AM, Isaacs N, Moretta M, Hicks GA, Urso DM, Brierley SM. Increased κ-opioid receptor expression and function during chronic visceral hypersensitivity. Gut 2014;63(7):1199–1200.

[38] Inserra MC, Israel MR, Caldwell A, Castro J, Deuis JR, Harrington AM, Keramidas A, Garcia-Caraballo S, Maddern J, Erickson A, Grundy L, Rychkov GY, Zimmermann K, Lewis RJ, Brierley SM, Vetter I. Multiple sodium channel isoforms mediate the pathological effects of Pacific ciguatoxin-1. Sci Rep 2017;7:42810.

[39] Isensee J, Krahe L, Moeller K, Pereira V, Sexton JE, Sun X, Emery E, Wood JN, Hucho T. Synergistic regulation of serotonin and opioid signaling contributes to pain insensitivity in Nav1.7 knockout mice. Sci Signal 2017;10(461).

[40] Israel MR, Tanaka BS, Castro J, Thongyoo P, Robinson SD, Zhao P, Deuis JR, Craik DJ, Durek T, Brierley SM, Waxman SG, Dib-Hajj SD, Vetter I. NaV 1.6 regulates excitability of mechanosensitive sensory neurons. J Physiol 2019;597(14):3751–3768.

[41] Jiang Y, Castro J, Blomster LV, Agwa AJ, Maddern J, Schober G, Herzig V, Chow CY, Cardoso FC, Demetrio De Souza Franca P, Gonzales J, Schroeder CI, Esche S, Reiner T, Brierley SM, King GF. Pharmacological Inhibition of the Voltage-Gated Sodium Channel NaV1.7 Alleviates Chronic Visceral Pain in a Rodent Model of Irritable Bowel Syndrome. ACS Pharmacol Transl Sci 2021;4(4):1362–1378.

[42] King GF, Vetter I. No gain, no pain: NaV1.7 as an analgesic target. ACS Chem Neurosci 2014;5(9):749–751.

[43] Koninckx PR, Ussia A, Adamyan L, Wattiez A, Gomel V, Martin DC. Pathogenesis of endometriosis: the genetic/epigenetic theory. Fertil Steril 2019;111(2):327–340.

[44] Kotecha M, Cheshire WP, Finnigan H, Giblin K, Naik H, Palmer J, Tate S, Zakrzewska JM. Design of Phase 3 Studies Evaluating Vixotrigine for Treatment of Trigeminal Neuralgia. J Pain Res 2020;13:1601–1609.

[45] Lee MC, Nahorski MS, Hockley JRF, Lu VB, Ison G, Pattison LA, Callejo G, Stouffer K, Fletcher E, Brown C, Drissi I, Wheeler D, Ernfors P, Menon D, Reimann F, St John Smith E, Woods CG. Human labour pain is influenced by the voltage-gated potassium channel K<sub>V</sub>6.4 subunit. bioRxiv 2020:489310.

[46] Li J, Stratton HJ, Lorca SA, Grace PM, Khanna R. Small molecule targeting NaV1.7 via inhibition of the CRMP2-Ubc9 interaction reduces pain in chronic constriction injury (CCI) rats. Channels (Austin) 2022;16(1):1–8.

[47] Maddern J, Grundy L, Castro J, Brierley SM. Pain in Endometriosis. Front Cell Neurosci 2020;14:590823.

[48] Maddern J, Grundy L, Harrington A, Schober G, Castro J, Brierley SM. A syngeneic inoculation mouse model of endometriosis that develops multiple comorbid visceral and cutaneous pain like behaviours. Pain 2022;163(8):1622–1635.

[49] Maertens C, Cuypers E, Amininasab M, Jalali A, Vatanpour H, Tytgat J. Potent modulation of the voltage-gated sodium channel Nav1.7 by OD1, a toxin from the scorpion Odonthobuthus doriae. Mol Pharmacol 2006;70(1):405–414.

[50] McAllister SL, McGinty KA, Resuehr D, Berkley KJ. Endometriosis-induced vaginal hyperalgesia in the rat: role of the ectopic growths and their innervation. Pain 2009;147(1-3):255–264.

[51] Michel MC, Igawa Y. Therapeutic targets for overactive bladder other than smooth muscle. Expert Opin Ther Targets 2015;19(5):687–705.

[52] Minett MS, Pereira V, Sikandar S, Matsuyama A, Lolignier S, Kanellopoulos AH, Mancini F, Iannetti GD, Bogdanov YD, Santana-Varela S, Millet Q, Baskozos G, MacAllister R, Cox JJ, Zhao J, Wood JN. Endogenous opioids contribute to insensitivity to pain in humans and mice lacking sodium channel Nav1.7. Nat Commun 2015;6:8967.

[53] Mueller A, Starobova H, Morgan M, Dekan Z, Cheneval O, Schroeder CI, Alewood PF, Deuis JR, Vetter I. Antiallodynic effects of the selective NaV1.7 inhibitor Pn3a in a mouse model of acute postsurgical pain: evidence for analgesic synergy with opioids and baclofen. Pain 2019;160(8):1766–1780.

[54] Muratspahic E, Tomasevic N, Koehbach J, Duerrauer L, Hadzic S, Castro J, Schober G, Sideromenos S, Clark RJ, Brierley SM, Craik DJ, Gruber CW. Design of a Stable Cyclic Peptide Analgesic Derived from Sunflower Seeds that Targets the kappa-Opioid Receptor for the Treatment of Chronic Abdominal Pain. J Med Chem 2021;64(13):9042–9055.

[55] Sadeghi M, Erickson A, Castro J, Deiteren A, Harrington AM, Grundy L, Adams DJ, Brierley SM. Contribution of membrane receptor signalling to chronic visceral pain. Int J Biochem Cell Biol 2018;98:10–23.

[56] Salvatierra J, Castro J, Erickson A, Li Q, Braz J, Gilchrist J, Grundy L, Rychkov GY, Deiteren A, Rais R, King GF, Slusher BS, Basbaum A, Pasricha PJ, Brierley SM, Bosmans F. NaV1.1 inhibition can reduce visceral hypersensitivity. JCI Insight 2018;3(11).

[57] Siebenga P, van Amerongen G, Hay JL, McDonnell A, Gorman D, Butt R, Groeneveld GJ. Lack of Detection of the Analgesic Properties of PF-05089771, a Selective Nav 1.7 Inhibitor, Using a Battery of Pain Models in Healthy Subjects. Clin Transl Sci 2020;13(2):318–324.

[58] Stratton P, Berkley KJ. Chronic pelvic pain and endometriosis: translational evidence of the relationship and implications. Hum Reprod Update 2011;17(3):327–346.

[59] Wang C, Wang Z, Yang Y, Zhu C, Wu G, Yu G, Jian T, Yang N, Shi H, Tang M, He Q, Lan L, Liu Q, Guan Y, Dong X, Duan J, Tang Z. Pirt contributes to uterine contraction-induced pain in mice. Mol Pain 2015;11:57.

[60] Zakrzewska JM, Palmer J, Morisset V, Giblin GM, Obermann M, Ettlin DA, Cruccu G, Bendtsen L, Estacion M, Derjean D, Waxman SG, Layton G, Gunn K, Tate S, study i. Safety and efficacy of a Nav1.7 selective sodium channel blocker in patients with trigeminal neuralgia: a double-blind, placebo-controlled, randomised withdrawal phase 2a trial. Lancet Neurol 2017;16(4):291–300.

[61] Zito G, Luppi S, Giolo E, Martinelli M, Venturin I, Di Lorenzo G, Ricci G. Medical treatments for endometriosis-associated pelvic pain. Biomed Res Int 2014;2014:191967.

[62] Zondervan KT, Becker CM, Koga K, Missmer SA, Taylor RN, Vigano P. Endometriosis. Nat Rev Dis Primers 2018;4(1):9.

